# Marine bacteria cross-feeding controls the fate of extracellular glycolate carbon

**DOI:** 10.1101/2025.09.29.679071

**Authors:** Ty J. Samo, Jeffrey A. Kimbrel, Kristina A. Rolison, Steven J. Blazewicz, Keith D. Morrison, Peter K. Weber, Xavier Mayali

## Abstract

Glycolate is a major product of phytoplankton photorespiration, but its fate in the microbial food web is not well constrained. Here, we used stable isotope probing and mass spectrometry combined with genomic analyses and microscopy to quantify glycolate metabolism by a taxonomically diverse set of heterotrophic marine bacteria. We found that 9 of 16 tested strains with the genomic capability to metabolize glycolate directly assimilated and respired glycolate carbon in monoculture. We next co-cultivated glycolate-incorporating strains with non-incorporating strains and found that several cross-feeders incorporated more glycolate carbon into their biomass than direct incorporators. Carbon use efficiency, reflecting proportional differences in movement of glycolate carbon into biomass versus into carbon dioxide, were distinct across co-cultures and ranged from 0.01 -3.15% depending on the strain mixtures. These results suggest that the fate of glycolate carbon is not limited to microbial taxa with the genetic capability for direct assimilation, and that bacterial metabolic interactions via cross-feeding play a critical role in influencing the efficiency of carbon transfer. Such information is critical to refine conceptual and numerical models of heterotrophic processing and transfer of organic carbon in an era of global change with predicted increases in photorespiration.

## Introduction

Glycolate (HOCH_2_CO_2_H; glycolic acid) is a simple organic molecule annually produced at the petagram scale primarily by photorespiration in autotrophs wherein ribulose-1,5-bisphosphate carboxylase/oxygenase (RuBisCo) binds oxygen (O_2_) instead of carbon dioxide (CO_2_) to generate phosphoglycolate instead of glycerate-3-phosphate[1–5]. In marine environments, photorespiration rate increases with increasing oxygen concentration [6] and light [7], leading to substantial glycolate production that reaches micromolar concentrations [8]. Turnover has been found to be around one week, although it can be as high as one hour [9]. Glycolate is therefore a central component of microbial food webs, especially in oligotrophic systems, where bacterial biomass and metabolism drive the biogeochemistry of the sunlit ocean [10]. It has been hypothesized that climate change may increase the flux of glycolate into the oceans due to nutrient and temperature stresses elevating rates of photosynthetic/photorespiratory overflow [11]. Thus, a better understanding of glycolate metabolism by marine bacteria is needed to understand how bacterial glycolate metabolism is partitioned into biomass and energy production, how bacterial interactions further influence these pathways, and how these dynamics influence C fate.

The substantial global pool of glycolate in seawater has prompted studies investigating its contribution to microbial biomass yield in natural systems and laboratory cultures [12], as well as its utilization by photoheterotrophs under varying light conditions [13]. New insights into the genome-encoded abilities of marine bacteria to metabolize glycolate continue to be discovered [14, 15], highlighting an opportunity to link bacterial taxonomy and genomic potential to glycolate processing and the fate of glycolate carbon. Three known transporters (glycolate permease, lactate permease, and the glycolate oxidase complex) bring the substrate into the cell where it is oxidized to glyoxylate (if not oxidized during transport) and then directed into central carbon metabolism via the glycerate pathway and/or the beta-hydroxyaspartate cycle (BHAC)[14]. Notably, glycolate assimilation through the glycerate pathway results in the release of CO_2_, whereas the BHAC incorporates all carbon into biomass without CO_2_ loss. This highlights how different bacterial genotypes can determine both the metabolic fate of glycolate and the overall fate of its carbon in the environment.

To interrogate how microbial identity relates to the path of carbon through a microbial food web, we used simplified culture and co-culture systems incubated with glycolate as the sole carbon source. First, we characterized the genomic capability of glycolate metabolism of the cultures, and we then empirically measured their glycolate incorporation and respiration. We then combined cultures that directly metabolized glycolate with those that did not to examine glycolate carbon cross-feeding. We sought to expand upon approaches that have employed carbon use efficiency (CUE; also known as bacterial growth efficiency or BGE) to understand C fluxes driven by microbial metabolism in bulk consortia [16–20] by examining this parameter in simplified cultures and laboratory settings. By measuring fluxes of glycolate carbon into biomass and CO_2_ in both pure heterotrophic cultures and co-cultures, we were able to account for taxonomic identity and activity of cross-feeding organisms to gain a more comprehensive understanding of glycolate metabolism in the microbial loop.

## Results

### 1 Comparative genomics revealed class-level patterns in glycolate metabolism

We leveraged our culture collection of sixteen heterotrophic bacteria isolated from cultures of the diatom *Phaeodactylum tricornutum* to examine how a subset of strains that rely on algal exudates contribute to glycolate C cycling. Thirteen isolates in this study are from phylum Pseudomonadota (eleven Alphaproteobacteria, one Gammaproteobacteria, and one Betaproteobacteria). The remaining three isolates belong to phylum Bacteroidota – two are Flavobacteriia and one is a Cytophagia (Fig. 1). All strains, except *Tepidicaulis* and *Yoonia*, have been previously reported in association with algal hosts [21–33], which are the primary source of glycolate in surface aquatic ecosystems. Further, querying the TARA Oceans dataset using Ocean Gene Atlas [34, 35] revealed that the 16S rRNA gene sequences of all strains were detected in the surface and deep chlorophyll maximum of coastal and open ocean regions across the world.

**Figure 1.**
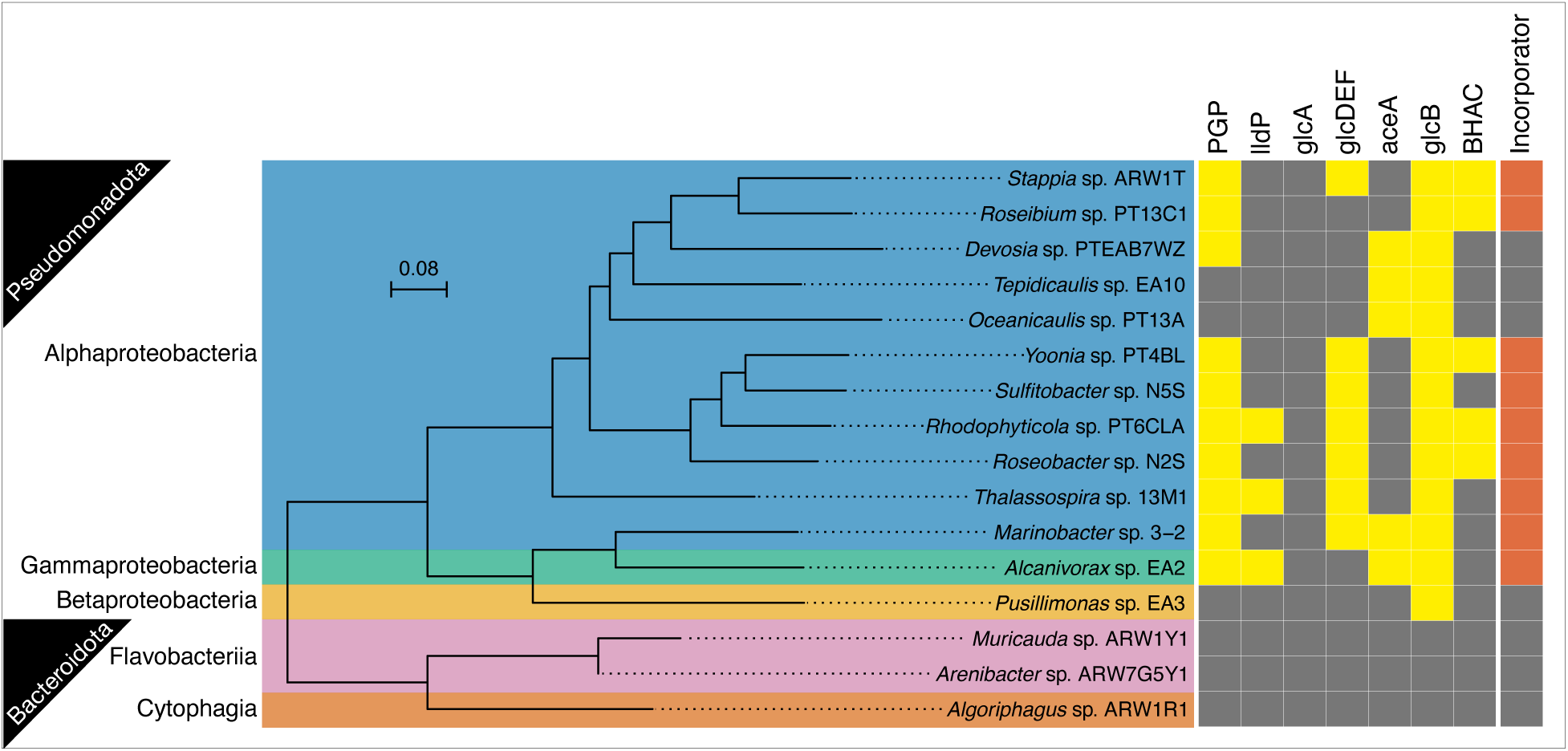
Genomic potential to assimilate glycolate aligns with phylogeny. Yellow indicates presence of the gene/pathway, and gray indicates absence. All Pseudomonadota contain at least one gene to metabolize glycolate while the Bacteroidota do not. Gene abbreviations: PGP, phosphoglycolate phosphatase; lldP, lactate permease (dehydrogenase); glcA glycolate permease; glcDEF, glycolate dehydrogenase/oxidase complex; aceA, isocitrate lyase; glcB, malate synthase; BHAC, B-hydroxyaspartate cycle. Incorporator indicates the strains were confirmed (orange) to assimilate glycolate C by isotope tracing and nanoSIMS in monoculture.

We examined the sequenced genomes of the sixteen isolates for the capability for glycolate metabolism across phyla and classes and to compare this potential to empirically measured incorporation rates. All Pseudomonadota possessed genes for glycolate metabolism. In contrast, the Bacteroidota did not contain glycolate metabolism genes, indicating no known capability for glycolate incorporation (Fig. 1). Examination at the class level revealed that there were no identified genes to specifically import glycolate via glycolate permease (glcA) in the eleven Alphaproteobacteria or the Gammaproteobacterium. Instead, there were several instances of the lactate permease gene (lldP), which is known to transport glycolate at the same rate as lactate [36, 37] and the glycolate dehydrogenase/oxidase complex (glcDEF), which enables glycolate import and conversion to glyoxylate in a single enzymatic process [38, 39]. Alphaproteobacteria was the only class to contain the BHAC genes for glycolate metabolism via the beta-hydroxyaspartate pathway [14]. The sole Betaproteobacterium contained one glycolate incorporation gene, malate synthase (glcB), which was also present in all the Pseudomonadota.

### 3 Stable Isotope Probing Corroborated Genome Prediction of Glycolate Metabolism

We compared genomic capacity for glycolate metabolism to functional activity with stable isotope probing (Fig. 2). Addition of ^13^C-glycolate as the sole carbon source to the monocultures enabled two independent methods to confirm genome-prediction of glycolate metabolism by quantifying its use for biomass (anabolism) and energy production (catabolism). We defined biomass production and energy production, respectively, as ^13^C-enrichment of single-cells and bulk CO_2_ using units of C_net_ d^-1^, the percent of biomass C or CO_2_ produced from glycolate per day [40, 41]. The three Bacteroidota strains with no genomic evidence for known glycolate metabolism did not incorporate C from glycolate into their biomass (Fig. 2A), nor respired glycolate C into CO_2_ (measured by cavity ring down spectroscopy, CRDS; Fig. 2B). The *Amylibacter* strain exhibited the highest glycolate C incorporation, obtaining 8% of its biomass from glycolate per day. On the other hand, the *Yoonia* culture respired the most glycolate C into CO_2_. Incorporation and respiration were linked: the same strains that exhibited C incorporation into biomass also produced CO_2_ from glycolate: *Thalassospira*, *Stappia*, *Amylibacter*, *Roseibium*, *Sulfitobacter*, *Alcanivorax*, *Marinobacter*, *Rhodophyticola*, and *Yoonia* (Fig. 2). Some of the other strains with low incorporation and/or respiration were statistically different from background but represent physiologically negligible activity and we considered the remaining strains as inactive: *Oceanicaulis*, *Muricauda*, *Pusillimonas*, *Devosia*, *Arenibacter*, *Algoriphagus*, and *Tepidicaulis*. Within strains that incorporated glycolate C, we observed cell-to-cell variability that spanned two orders of magnitude (Fig. 2a), consistent with reports in other isogenic microbial systems [42]. Respiration of glycolate largely matched the presence of glycolate metabolism genes in all but four genera - *Devosia*, *Tepedicaulis*, *Oceanicaulis*, and *Pusillimonas* (Fig. 1 and 2B) – that exhibited glycolate metabolism genes but no detectable respiration or assimilation. These results thus differentiated glycolate metabolism pathways that were responsible for uptake of extracellular glycolate into the cells versus conversion/regeneration of intracellular pools (e.g. phosphoglycolate generated from DNA repair)[43] and confirmed that a strain’s lack of any glycolate genes corresponded with no uptake.

**Figure 2.**
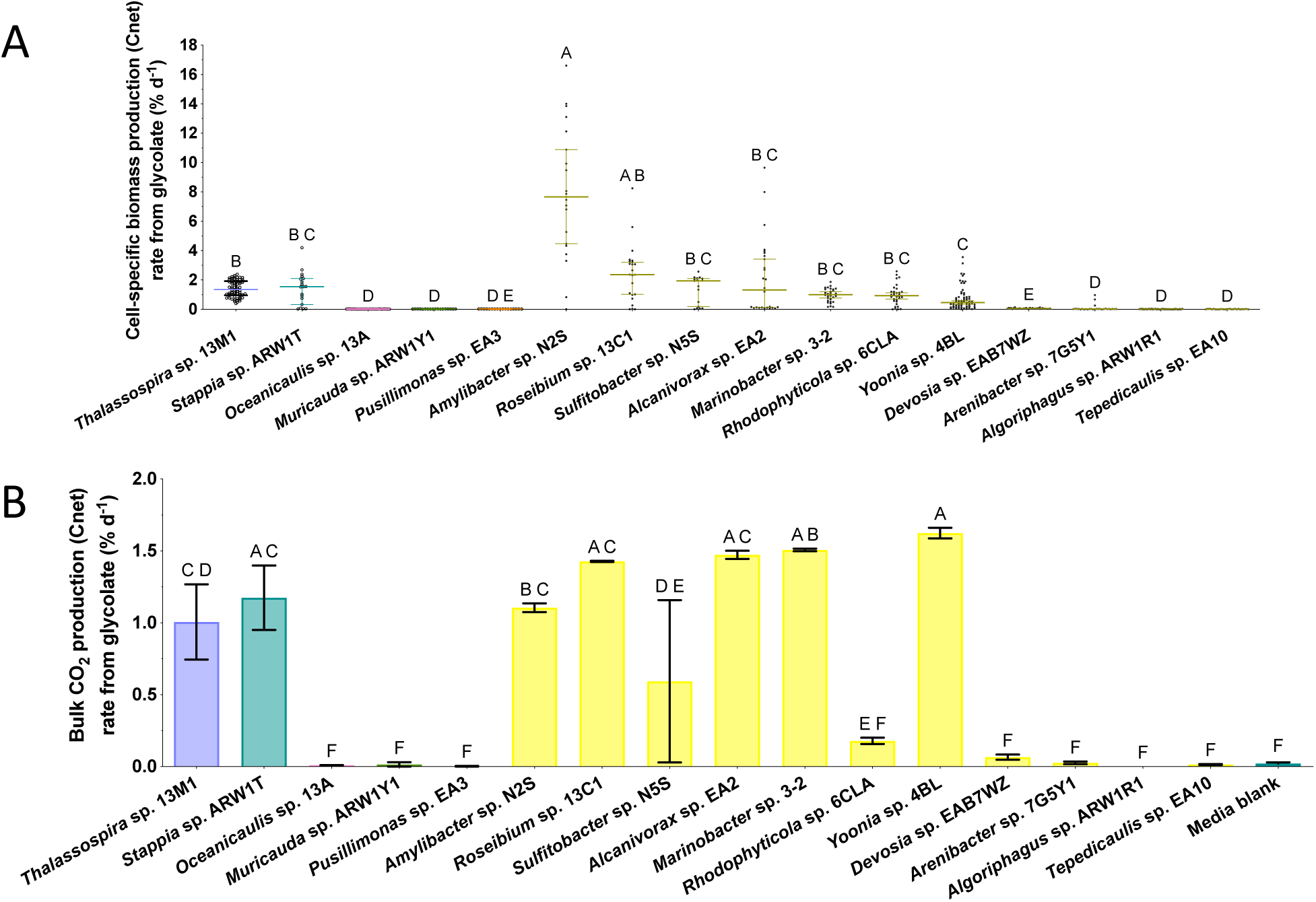
Stable isotope probing of ^13^C from glycolate, measured as C_net_ % d,^-1^ within individual cells (A) and in CO_2_ (B) provided direct evidence of bacterial isolates’ ablilities to metabolize glycolate. Letters above plots denote statistical differences with shared letters indicating no statistical difference between median values in single-cell nanoSIP data (Kruskal-Wallis tests) and mean values in the CO_2_ data (one-way ANOVA). For subsequent co-cultivation experiments, we selected the 5 isolates on the left of the plots: Thalassospira and Stappia as incorporators and Oceanicaulis, Muricauda, and Pusillimonas as non-incorporators.

### 4 Selection of incorporators and non-incorporators to examine carbon use efficiency of co-cultures

Out of the nine strains that assimilated glycolate (hereafter “incorporators”), we selected two (*Thalassospira* and *Stappia*) to assess whether their broad ecophysiological similarities translated into similar interactions with strains that did not assimilate glycolate. Both incorporators are Alphaproteobacteria globally distributed throughout the ocean’s water column (surface, deep chlorophyll maximum, and mesopelagic zones [34, 35]), exhibit a free-living lifestyle when cultivated with the diatom *Phaeodactylum tricornutum* [40], and had statistically equivalent percentages of their biomass produced from glycolate (Mann-Whitney test, p = 0.99; Figure 2A). Out of the seven strains that did not assimilate glycolate (hereafter “non-incorporators”), we selected three that are taxonomically diverse and globally distributed: *Oceanicaulis*, *Muricauda*, and *Pusillimonas*. We aimed to compare how their taxonomy and genomic capacities to metabolize glycolate related to their uptake of glycolate-derived compounds released by *Thalassospira* and *Stappia*.

The genomes of the three chosen non-incorporators exhibited different potential for the metabolism of glycolate-derived metabolites. *Oceanicaulis* contains the malate synthase gene (glcB), which converts glycolate or its oxidized form, glyoxylate, to malate, but no additional glycolate metabolism genes. This limited capacity was consistent with the observed lack of glycolate uptake in monoculture, suggesting that *Oceanicaulis* primarily recycled its own glycolate pools, such as those converted from central carbon substrates via isocitrate lyase (aceA), which converts isocitrate to glyoxylate [44]. In contrast to *Oceanicaulis*, *Pusillimonas* (Gammaproteobacteria) only contained glcB and no aceA, suggesting more restricted glycolate metabolism. Amino acids such as glycine [45] can be processed to generate glyoxylate that would enter central carbon metabolism via glcB. The third non-incorporator, *Muricauda* (Bacteroidota), lacked any genes associated with glycolate metabolism, representing an absent glycolate metabolism phenotype characterized by an inability to transport glycolate into the cell, oxidize it to glyoxylate, and ultimately perform the glyoxylate shunt, which converts glycolate/glyoxylate to glucose [46].

After incubating the six co-cultures for 7 days separated by a membrane that allows diffusion of metabolites but no physical attachment, glycolate was not detectable in either side of the vessels (below 500 nM), indicating that > 99% of the 50 µM added glycolate was assimilated into biomass, respired, or converted to other metabolites and excreted (excretedFig. S2). We also measured the ^13^C composition of the DIC from each side of the incubation bottles to quantify respiration of glycolate, as done in the monocultures. Using NanoSIMS, we quantified the uptake of glycolate C by the incorporator and non-incorporator cells. Unlike in monoculture where assimilation was not detected, *Oceanicaulis* incorporated statistically greater glycolate C than formalin-killed controls when co-cultured with incorporators *Thalassospira* (Fig. 3A) or *Stappia* (Fig. 4A). In co-culture with *Thalassospira*, *Oceanicaulis* assimilation was relatively low (median 0.03% d^-1^). In co-culture with *Stappia*, most *Oceanicaulis* cells did not incorporate glycolate derived C (median 0.005% d^-1^), but a subset of the cells incorporated as much C as the *Stappia* incorporators (maximum 2.6% d^-1^).

**Figure 3.**
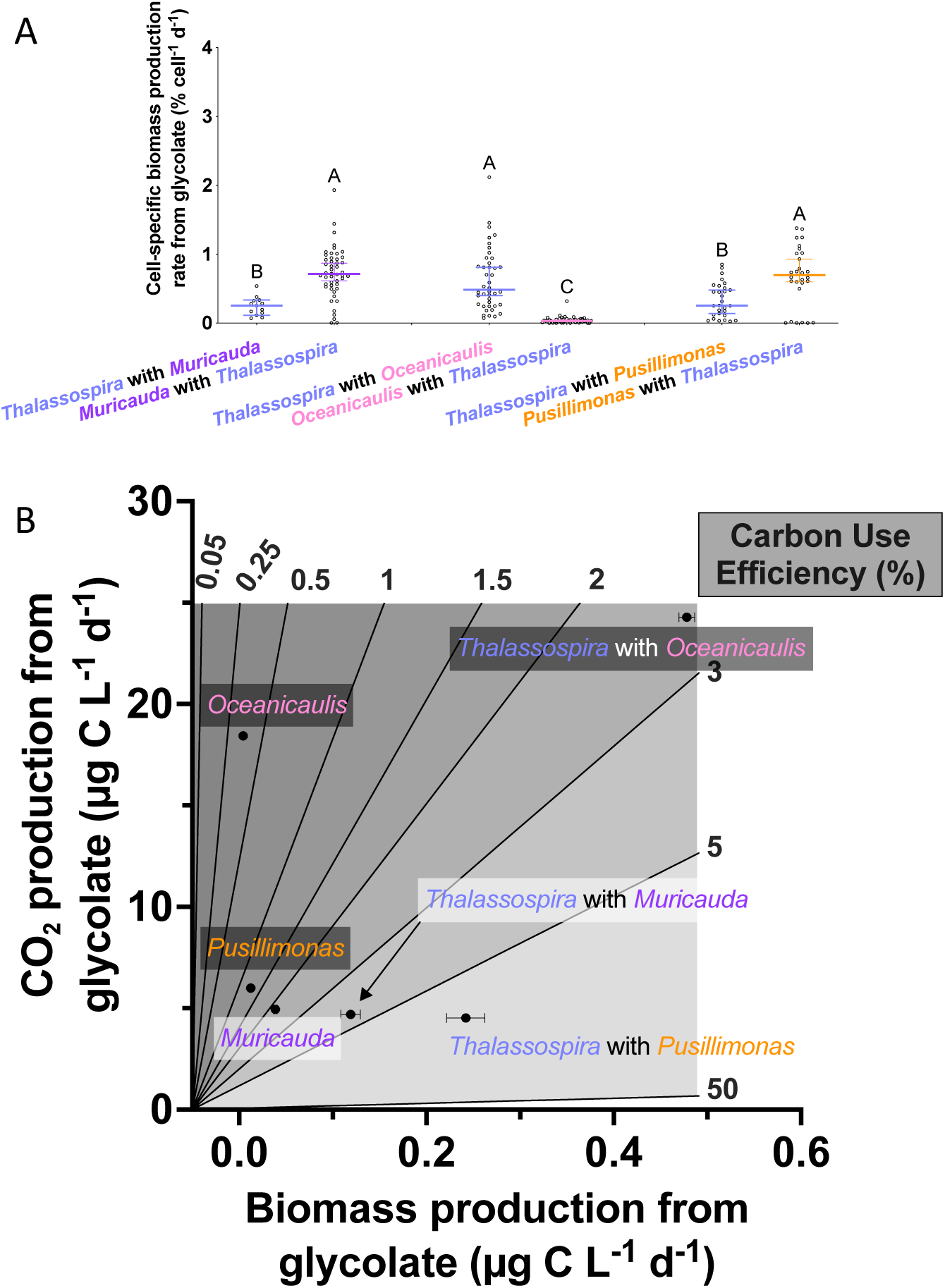
Cell-specific biomass production (A) and bulk carbon use efficiency, biomass production, and respiration (B) from glycolate or glycolate-derived metabolites of primary incorporator Thalassospira and co-culture partners Muricauda, Oceanicaulis, and Pusillimonas. At the single-cell level, non-incorporators Muricauda and Pusillimonas generated as much or greater biomass from glycolate-derived compounds as the incorporators did from glycolate. Thalassospira CUE was greatest when co-cultured with Pusillimonas, while its biomass and CO_2_ production was greatest when co-cultured with Oceanicaulis. Meanwhile, Muricauda CUE of glycolate-derived metabolites from Thalassospira is significantly greater than Oceanicaulis but not statistically distinguishable from Pusillimonas. Similarly, Muricauda also exhibited greater biomass production from secondary metabolites than Oceanicaulis but not Pusillimonas. CO_2_ production was greatest in the Oceanicaulis, followed by Pusillimonas, and then Muricauda. Statistical groupings based on ANOVAs for panel B are noted here: CUE: Thalassospira w/ Pusillimonas > Thalassospira w/ Muricauda = Thalassospira w/ Oceanicaulis > Muricauda, Pusillimonas > Pusillimonas, Oceanicaulis Production: Thalassospira w/ Oceanicaulis > Thalassospira w/ Pusillimonas > Thalassospira w/ Muricauda > Muricauda, Pusillimonas > Pusillimonas, Oceanicaulis Respiration: Thalassospira w/ Oceanicaulis > Oceanicaulis > Pusillimonas > Muricauda > Thalassospira w/ Muricauda > Thalassospira w/ Pusillimonas

Unlike *Oceanicaulis*, the other two non-incorporators showed a distinct response when co-cultivated with glycolate incorporators. Non-incorporator *Muricauda* incorporated glycolate C when co-cultured either with *Thalassospira* (Fig. 3A) or with *Stappia* (Fig. 4A). In both cases, *Muricauda* cells incorporated more C from glycolate as a fraction of their biomass (C_net_) than the direct incorporators *Thalassospira* and *Stappia*. Non-incorporator *Pusillimonas* did not incorporate glycolate C in co-culture with *Stappia* (Fig. 4A), but incorporated glycolate C when co-cultured with *Thalassospira* (Fig. 3A). Here again at the single cell level, *Pusillimonas* incorporated more of its biomass C from glycolate than *Thalassospira* cells. These results suggest that glycolate incorporators *Stappia* and *Thalassospira* released glycolate-derived C into metabolites that subsequently became available to some (but not all) of the non-incorporators. Further, the non-incorporators were relatively efficient in the assimilation of the glycolate-derived metabolites (except *Oceanicaulis* in co-culture with *Thalassospira*), as they took up relatively larger fractions of their biomass than the direct glycolate incorporators. While we detected significant quantities of ^13^C in biomass or CO_2_ in all non-incorporator co-cultures, supporting cross-feeding as one mechanism that shuttled C from glycolate (Figs. 3, 4, 5), we cannot rule out that the presence of the incorporator initiated regulatory changes and/or co-metabolism in the non-incorporator to enable uptake..

**Figure S2.**
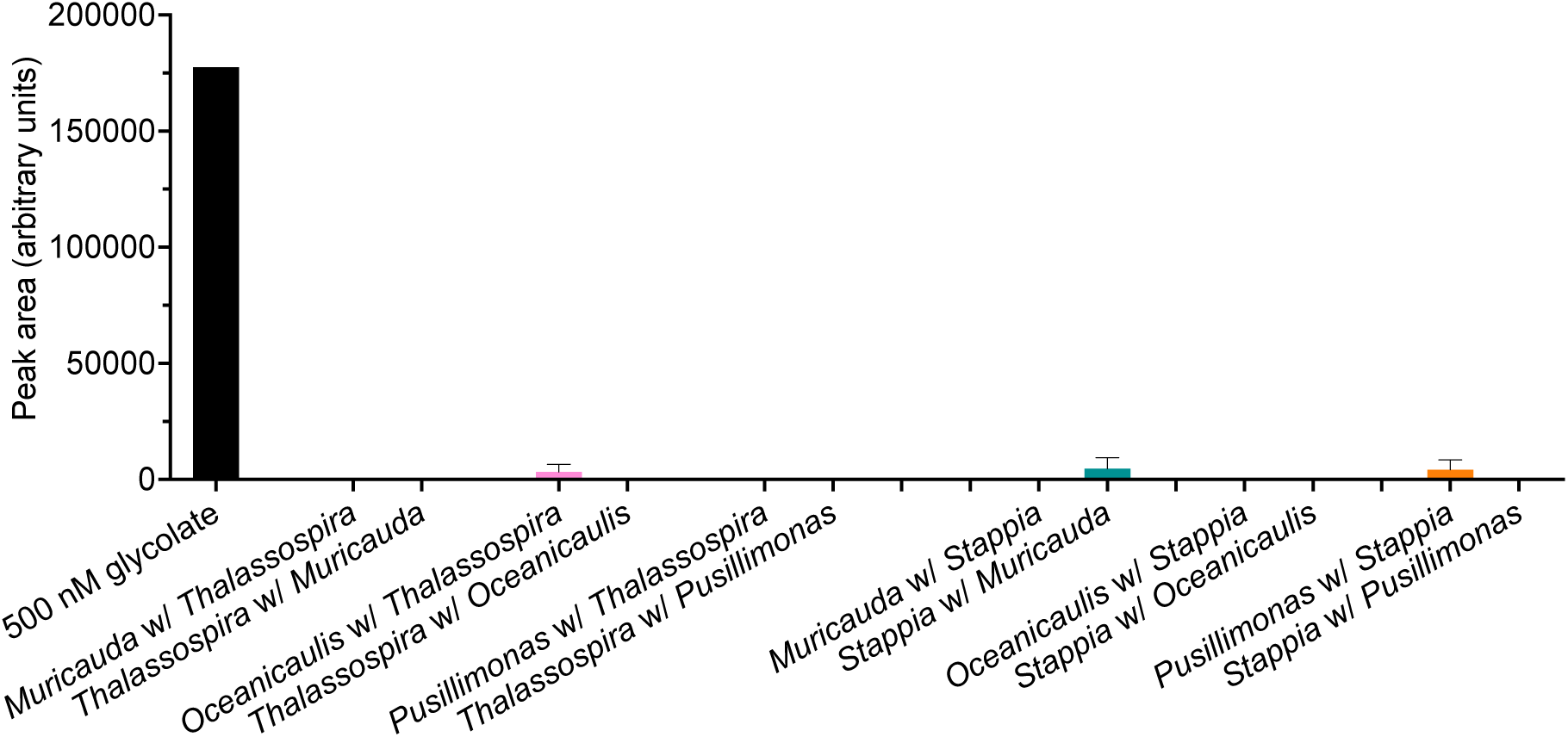
Removal of glycolate from the cultures demonstrated by peak areas of zero or lower than the 500 nm glycolate standard.

### 5 Total C flux from glycolate into *Thalassospira* and *Stappia* was affected by co-culture partner

Using the biomass C production rate from glycolate (measured by nanoSIMS), cell abundances, cell biovolumes, and bulk glycolate C respired as CO_2_ (measured by CRDS), we calculated total volumetric glycolate C assimilated and respired, allowing us to measure the taxon specific CUE (Fig. 3B, 4B) for co-cultures. Patterns of ^13^C-based glycolate CUEs in *Thalassospira* and *Stappia* showed that cultivation alongside non-incorporating partners impacted the balance between assimilation and respiration. For *Thalassospira*, CUE was significantly higher when co-cultivated with *Pusillimonas* (compared to *Muricauda* and *Oceanicaulis*, which were not significantly different from each other (Fig. 3B), leading us to reject the null hypothesis that CUE would not be impacted when cultivated alongside non-incorporating partners. This increase in efficiency was driven by proportionally more retention of glycolate C in biomass compared to when co-cultured with *Muricauda* and proportionally less glycolate C respired to CO_2_ compared to grown with *Oceanicaulis* (Fig. S2A). Further, we found that the percent of individual *Thalassospira* cells incorporating glycolate was significantly reduced in the *Muricauda* co-culture compared to the 100% cell incorporation observed with *Pusillimonas* (Table 1). Even-though cell specific C incorporation was higher for non-incorporators *Muricauda* and *Pusillimonas*, when taking into account total C (using cell biovolume and abundances), biomass production by *Thalassospira* was greater than the non-incorporator partners (Fig. 3B). Like Thalassospira, we rejected the null hypothesis that non-incorporators would not drive changes in *Stappia* CUE. However, the responses were distinct from those measured in *Thalassospira* and further supported how the identities of bacteria-bacteria partners influenced glycolate fate (Fig. 4B). *Stappia* CUE was highest when co-cultured with

**Table 1.** Percentages of glycolate-incorporating cells as influenced by non-incorporating partner.

| Culture | Co-culture partner | Glycolate assimilation (%) | s.e. | Statistical differences |
| --- | --- | --- | --- | --- |
| *Thalassospira* | *Muricauda* | 81.1 | 8.7 | A |
|  | *Oceanicaulis* | 99.2 | 1.7 | A B |
|  | *Pusillimonas* | 100 | 0 | B |
| *Stappia* | *Muricauda* | 96.0 | 4.5 | A |
|  | *Oceanicaulis* | 96.3 | 3.2 | A |
|  | *Pusillimonas* | 91.3 | 1.5 | A |

**Figure 5.**
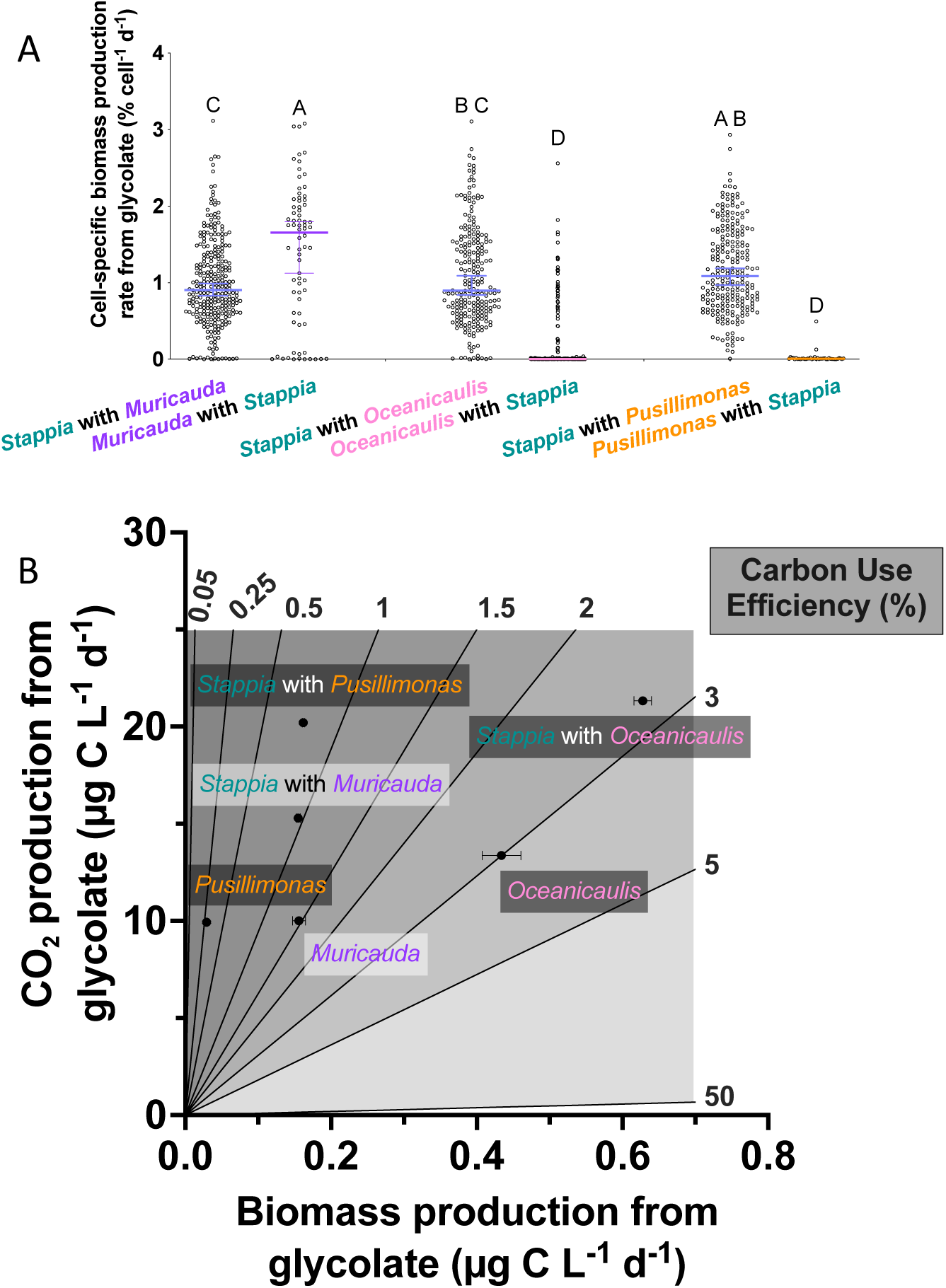
Cell-specific biomass production (A) and bulk carbon use efficiency, biomass production, and respiration (B) from glycolate or glycolate-derived metabolites of Stappia and co-culture partners Muricauda, Oceanicaulis, and Pusillimonas. Single-cells of non-incorporator Muricauda generated greater biomass from glycolate-derived compounds as Stappia did from glycolate while Oceanicaulis and Pusillimonas assimilated much less. Stappia CUE, biomass production, and respiration was greatest when co-cultured with Oceanicaulis. Oceanicaulis CUE of glycolate-derived metabolites from Stappia was significantly greater than all others, including its Stappia partner. Oceanicaulis also exhibited the greatest biomass production and respiration rates of the non-incorporators. Statistical groupings based on ANOVAs for panel B are noted here: CUE: Oceanicaulis > Stappia w/ Oceanicaulis > Muricauda > Stappia w/ Muricauda = Stappia w/ Pusillimonas > Pusillimonas Production: Stappia w/ Oceanicaulis > Oceanicaulis > Stappia w/ Muricauda = Stappia w/ Pusillimonas = Muricauda > Pusillimonas Respiration: Stappia w/ Oceanicaulis > Stappia w/ Pusillimonas > Stappia w/ Muricauda > Oceanicaulis > Muricauda = Pusillimonas

*Oceanicaulis*, driven by more retention of glycolate C into its biomass (Fig. S2B), and not significantly different when co-cultured with *Muricauda versus Pusillimonas*. These differences were not driven by changes in the fraction of *Stappia* cells with detectable incorporation (Table 1), indicating no major disruption in *Stappia*’s ability to assimilate glycolate by the non-incorporating partners. We found that CUE generally correlated with biomass and CO_2_ production values (Fig. 4B). Cellular biomass production from glycolate by *Thalassospira* and *Stappia* (C_net_) was always highest when *Oceanicaulis* was the non-incorporating partner. *Thalassospira* CO_2_ production from glycolate was also highest when partnered with *Oceanicaulis* compared to *Muricauda* and *Pusillimonas* (Fig. 3B).

**Figure S3.**
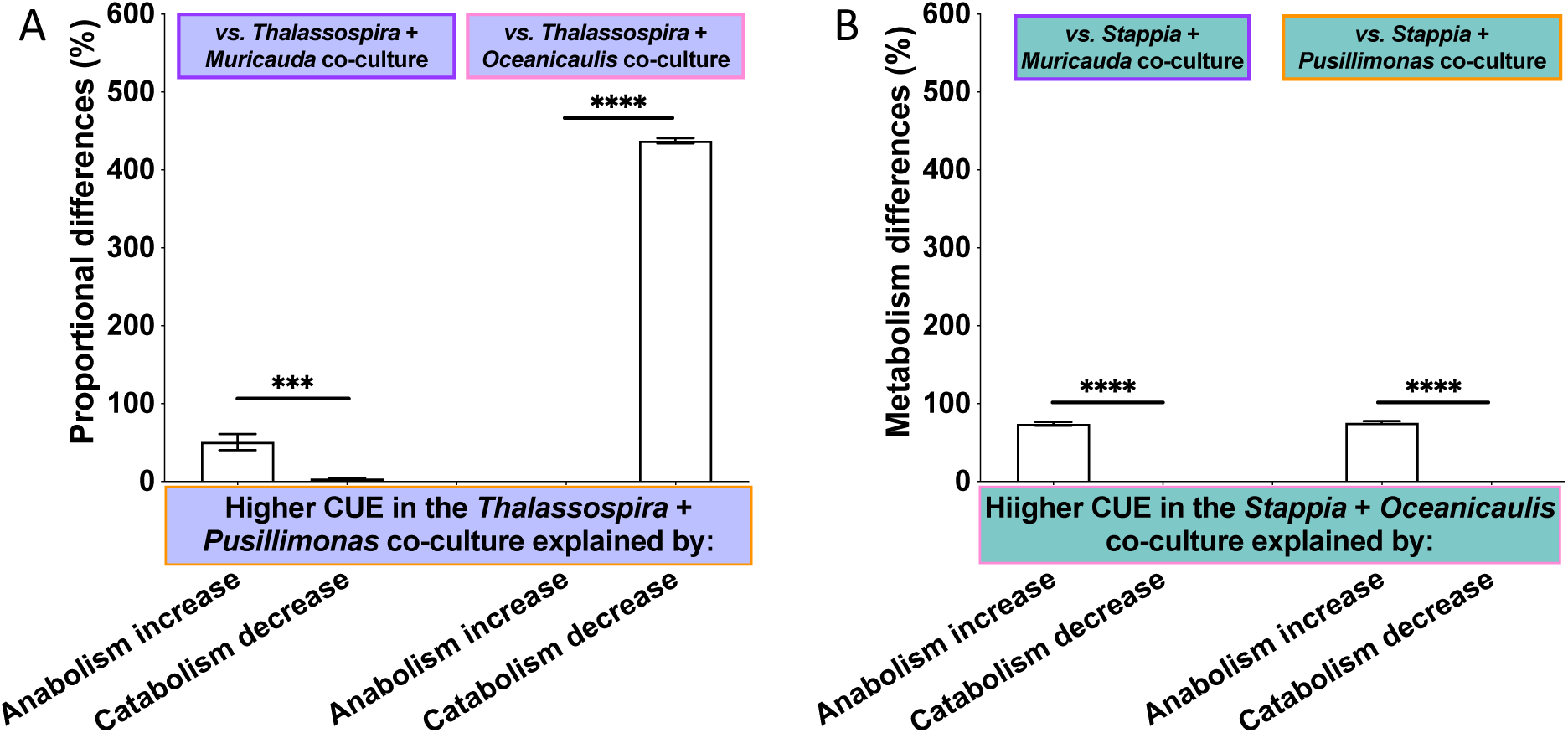
Proportional changes in anabolism (biomass production) versus catabolism (CO_2_ production) driving enhanced CUE in the Thalassospira + Pusillimonas (A) and Stappia + Oceanicaulis (B) co-cultures. Thalassospira exhibits higher CUE when co-cultured with Pusillimonas due to increased retention of glycolate into biomass compared to the Muricauda co-culture and decreased release of CO2 compared to the Oceanicaulis co-culture. Meanwhile, Stappia exhibits higher CUE when co-cultured with Oceanicaulis due to increased retention of glycolate into biomass compared to both the Muricauda and Pusillimonas co-cultures.

### 6 *Stappia* increased flux of glycolate carbon to cross-feeders compared to *Thalassospira*

The cross-feeding hypothesis was most tightly constrained for *Muricauda* due to its lack of annotated glycolate metabolism genes (this study), noted glycan processing activities [47], and preference for polymeric substrates [48]. We detected ^13^C in *Muricauda* biomass and headspace CO_2_, showing that ^13^C-substrates released from both incorporators were labile compounds readily metabolized by *Muricauda,* an organism with no known glycolate incorporation pathways. Secondary metabolites released by *Stappia* generated higher CUE and rates of anabolism and catabolism in *Muricauda* compared to when released by *Thalassospira* (Fig. 5). *Oceanicaulis* metabolism of glycolate-derived substrates diverged when co-cultivated with *Thalassospira* versus *Stappia* and supported release of more assimilable glycolate-derived compounds from *Stappia*. In addition to higher CUE when cultivated with *Stappia*, we found that *Oceanicaulis* biomass production rate from ^13^C-substrates was higher as well compared to *Thalassospira* (Fig. 5A), but the %-labeled cells were significantly lower with *Stappia* (Table 2, t-test, p < 0.05). In other words, there was a greater percentage of *Oceanicaulis* cells incorporating a small portion of *Thalassospira* secondary metabolites and a smaller percentage assimilating more metabolites from *Stappia*. This observation was met with significantly higher in ^13^CO_2_ production rate with *Thalassospira* compared to with *Stappia* (Fig. 5B). *Pusillimonas* exhibited low CUE regardless of partner, suggesting there was little difference in the preference of *Pusillimonas* for glycolate-derived metabolites released by either incorporator (Fig. 5C). We could not statistically distinguish CUE, biomass production, or %-labeling in *Pusillimonas* co-cultivated with *Thalassospira* versus *Stappia* (Fig. 5AB and Table 2), but CO_2_ production and cell C content were significantly higher with *Stappia* (Fig. 5B and S3).

Based on these results, we fully rejected the following two null hypotheses for the non-incorporator *Muricauda* and conditionally for *Oceanicaulis* and *Pusillimonas*: 1) There would be no CUE differences between non-incorporating strains when cultivated with the same glycolate-incorporating partner and 2) There would be no differences in the CUE between the same non-incorporating strains when cultivated with *Thalassospira* versus *Stappia*. CUE was not different between *Oceanicaulis* and *Pusillimonas* when co-cultured with *Thalassospira* and *Pusillimonas* CUE was indistinguishable between partnerships with *Thalassospira* versus *Stappia*.

## Discussion

### 1 Genomic and physiological insights into glycolate metabolism

Our observations of phylogenetic distinctions in glycolate metabolism highlighted how the presence or absence of bacterial taxa within a consortium can influence processing of specific substrates and control C biogeochemistry. While nearly all the isolates are noted members of algal microbiomes, Alphaproteobacteria appear to be the primary beneficiaries to photorespiration processes that increase availability of exogenous glycolate. Once incorporated, glycolate is generally oxidized to glyoxylate to fuel the glyoxylate shunt{Citation}, which efficiently replenishes intermediates of the TCA cycle to support anabolic and catabolic processes without generating two CO_2_ molecules [49]. The ability to metabolize C2-compounds via the glyoxylate cycle or larger compounds via the TCA cycle confers some plasticity in Alphaproteobacteria carbon metabolism and enables efficient assimilation of diverse substrates. This influence over carbon fate, combined with unique abilities to process other highly abundant components of marine DOM, may explain the widespread abundance of Alphaproteobacteria in surface oceans [50]. Moreover, Alphaproteobacteria may further impact DOM composition through the eventual exudation of substrates generated from glycolate to support cell wall production [51], metals acquisition [52], and oxidative stress responses [53].

We found that the presence of either glycolate dehydrogenase/oxidase complex (glcDEF) or lactate permease (lldP) was a prerequisite for direct assimilation of glycolate into biomass or respiration (Figs. 1 and 2). *Roseibium* was the notable exception, exhibiting uptake and respiration without these genes. However, the *Roseibium* genome contains genes for the BHAC pathway, which suggests the existence of yet-to-be-discovered glycolate transporter genes associated with this glycolate incorporation pathway. Meanwhile, the absence of glcDEF, lldP, and BHAC genes explained why *Devosia*, *Tepidicaulis*, *Oceanicaulis*, and *Pusillimonas* did not exhibit detectable assimilation despite having genes associated with glycolate metabolism. This was especially apparent for *Pusillimonas*, which only contained the malate synthase gene (glcB) that converts glyoxylate into malate with the help of acetyl-CoA. In these examples, the perceived mismatch between genome potential and observed physiology, as seen in other marine bacterial genera [40, 54, 55], is explained by whether the glycolate enzyme properties include transport into the cells. The inability to transport glycolate in these genera alludes to their potential occupation of specific biogeochemical niches via uptake and metabolism of alternative organic compounds that are converted to glycolate/glyoxylate intracellularly [56, 57]. Bacterial glycolate metabolism thus falls along a spectrum with distinctions driven by the unique nutrient requirements and genome profiles of each strain. In the absence of diverse substrates able to support central carbon metabolism, they (and other members like them) may thus rely on the excreted products of co-occurring taxa for subsistence or growth.

**Figure 8.**
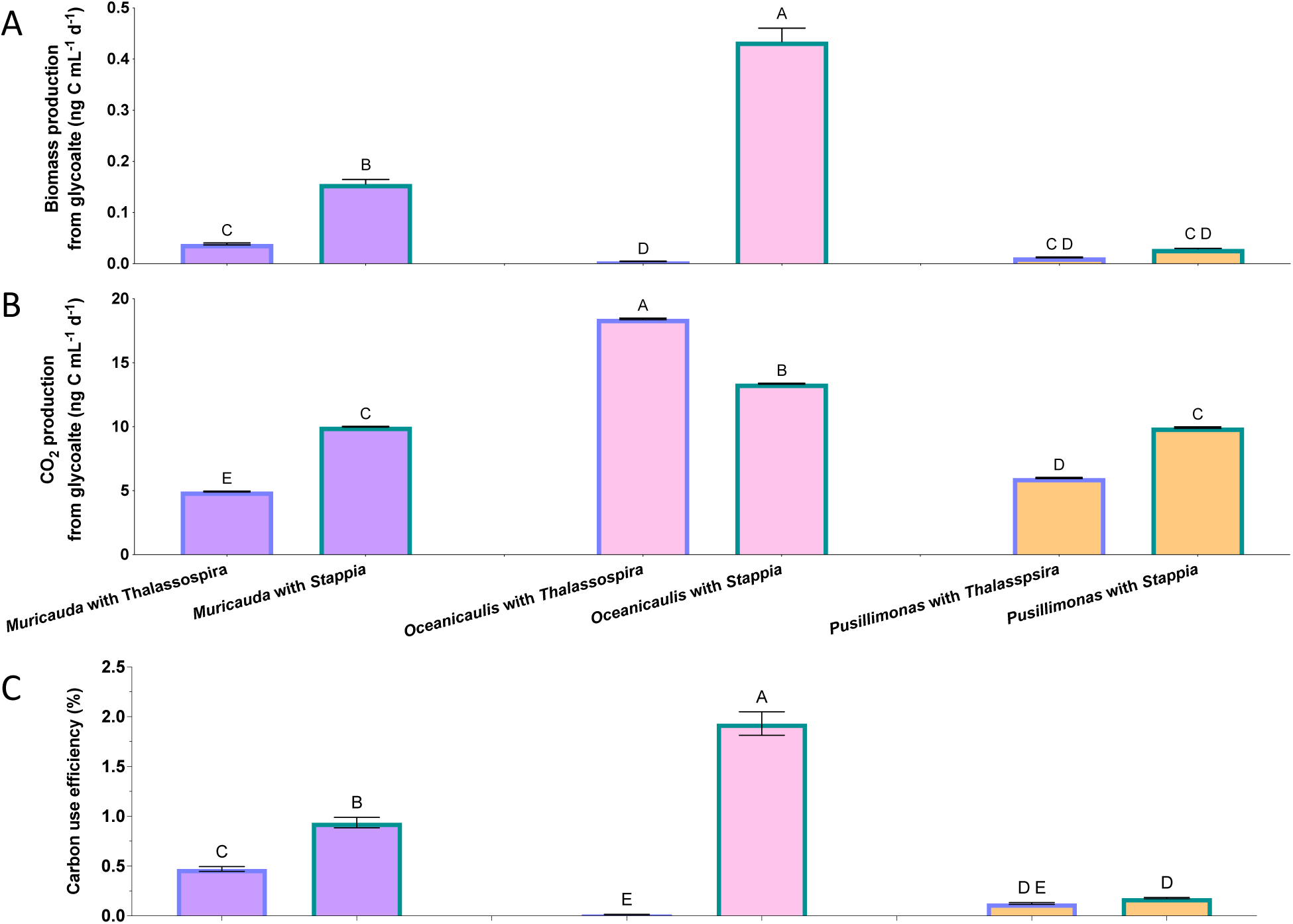
Comparison of CUE, biomass production, and CO_2_ production between non-incorporators reveals highest CUE and production when cultivated with Stappia compared to Thalassospira. Oceanicaulis featured the greatest biomass and CO_2_ production while Pusillimonas was consistently exhibited the lowest.

**Table 17.** Percentages of cells incorporating secondary metabolites generated from glycolate incorporators.

| Culture | Co-culture partner | Assimilation (%) | s.e. | Statistical differences |
| --- | --- | --- | --- | --- |
| *Muricauda* | *Thalassospira* | 78.1 | 7.0 | ns |
|  | *Stappia* | 50.1 | 16.9 |  |
| *Oceanicaulis* | *Thalassospira* | 76.3 | 3.8 | * |
|  | *Stappia* | 21.7 | 11.4 |  |
| *Pusillimonas* | *Thalassospira* | 31.6 | 30.0 | ns |
|  | *Stappia* | 13.7 | 4.9 |  |

The lack of notable glycolate metabolism within the Bacteroidota was consistent with their alternative lifestyles. As transformers of sporadic inputs of high molecular weight compounds [47, 58], these phyla rely on their associations with algal blooms and particulate organic matter (POM) to support their growth and energy needs. Their persistence in oligotrophic ocean microbiomes suggests that, in addition to sporulation [59, 60] or dormancy [61], they may rely on incorporation of other labile substances, and/or, like the non-incorporating Pseudomonadota above, assimilate secondary metabolites from primary incorporators to support their populations until the next encounter with blooms and/or high POM events.

### 2 Glycolate carbon use efficiencies in *Thalassospira* and *Stappia* were enhanced by different co-culture partners, but biomass production was enhanced by *Oceanicaulis*

#### Thalassospira

The influences of non-incorporator presence on *Thalassospira* glycolate metabolism revealed possible roles of non-incorporator physiology in modulating fate of glycolate C. The enhancement of *Thalassospira* CUE when co-cultured with *Pusillimonas* aligns with other reports on *Pusillimonas* participation in cross-feeding of degraded bisphenol A products [61]. Considering this phenotype alongside notable roles as an efficient degrader, reformer [62], and incorporator [50] of dissolved organic matter, *Pusillimonas* may have assimilated glycolate-based metabolites and then released tertiary metabolites that were preferentially incorporated by *Thalassospira* (i.e., tertiary metabolites or double cross-feeding). Quorum sensing [62] is another manner through which *Pusillimonas* may have shifted *Thalassospira* gene expression and metabolism, but the lack of studies examining chemical communication in *Pusillimonas* highlights a research gap and unconstrained mechanism in this system.

In the case of *Thalassospira* co-cultured with *Muricauda*, release of inhibitory compounds by *Muricauda* may have disrupted glycolate metabolism in a large proportion of the *Thalassospira* cell population. There are literature reports of *Muricauda* spp. that negatively impacted other Pseudomonads [63], but further studies are needed to determine the extent of this antimicrobial ability across *Muricauda* species. Another non-mutually exclusive explanation may be that *Muricauda* released unlabeled C in metabolites originating from having previously grown in high nutrient ZoBell media and then were preferentially assimilated by *Thalassospira*. This would add unlabeled C to the biomass and decrease ^13^C-enrichment. We observed significantly decreased biomass production from glycolate yet increased single-cell C content in this *Thalassospira* co-culture compared to the others, indicative of increased availability of unlabeled nutrients [64] via *Muricauda* but also consistent with disruption in cell division [65] (Fig. S4).

**Figure S4.**
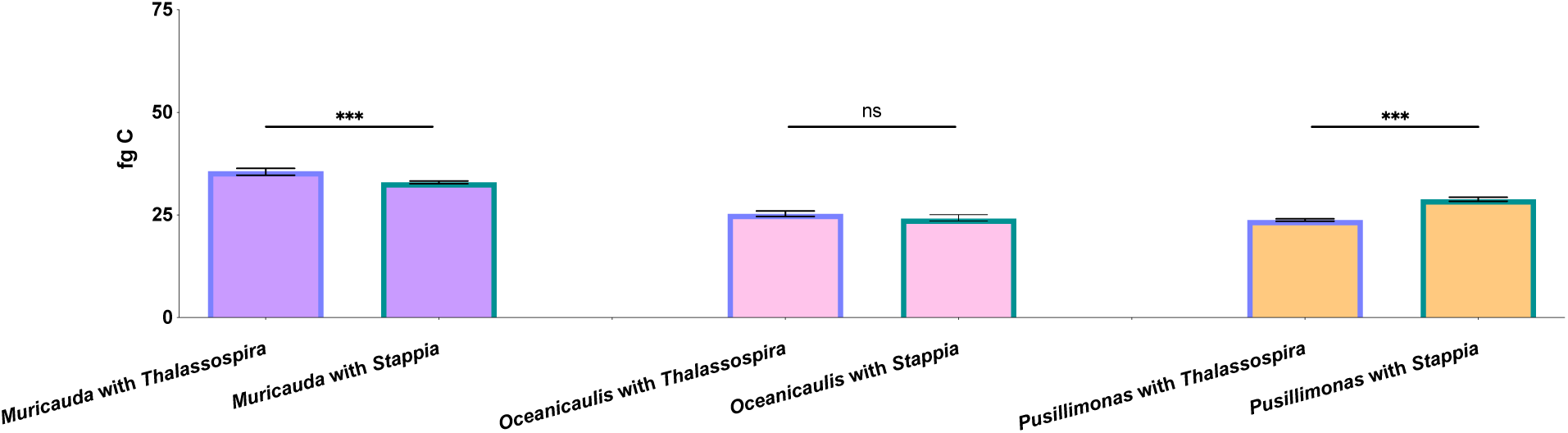
Median carbon content of non-incorporators cultivated with Thalassospira vs. Stappia as the incorporating partner.

Similarly, *Thalassospira* glycolate CUE was significantly reduced when cultivated with *Oceanicaulis* compared to cultivation with *Pusillimonas*. Like *Pusillimonas*, *Oceanicaulis* is a noted remineralizer of algal exudates [40] and may be have processed *Thalassospira*-released ^13^C-macromolecules and/or unlabeled macromolecules from previous growth in high nutrient media. However, a major difference is that the ^13^C re-incorporated by *Thalassospira* supported respiration processes, thus decreasing CUE. Unlabeled C-substrates could, as discussed in the *Muricauda* section above, dilute intracellular ^13^C pools and appear to reduce CUE. Uniquely however, *Thalassospira* exhibited the lowest single-cell C content (Fig. S4) but also highest rates of biomass and CO_2_ production (Fig. 3B), indicating that cells were likely nutrient stressed [66] but also rapidly metabolizing glycolate. Together, these observations suggest that *Oceanicaulis* impacted *Thalassospira* such that more total C from glycolate was metabolized and led to proportionately more CO_2_ release.

#### Stappia

The enhancement of *Stappia* CUE when co-cultivated with *Oceanicaulis*, relative to its partnerships with *Muricauda* and *Pusillimonas*, revealed an interaction that led to more biomass production from glycolate C. This observation contrasted with the reduced *Thalassospira* CUE when co-cultured with *Oceanicaulis* and suggested that exchanges of C between *Stappia* and *Oceanicaulis* were more likely to support biomass production. Like the *Thalassospira* co-culture, however, the presence of *Oceanicaulis* appeared to drive the highest rates of biomass and CO_2_ production compared to the presence of *Muricauda* and *Pusillimonas* (Fig. 4B), in addition to the highest median cell carbon content (Fig. S5). In this partnership, *Oceanicaulis* processing of glycolate-derived substrates from *Stappia* (i.e., tertiary metabolites) appears to have supported proportionately more anabolism compared to the *Thalassospira*-*Oceanicaulis* system. One explanation is an increase in substrate quality, which can drive higher CUE [67]; compounds requiring multiple enzymatic steps generate more CO_2_ and compounds with low degrees of reduction yield less energy. Cross-feeding of higher quality substrates produced from an individual precursor may thus be an influential component affecting bacterial CUE.

**Figure S5.**
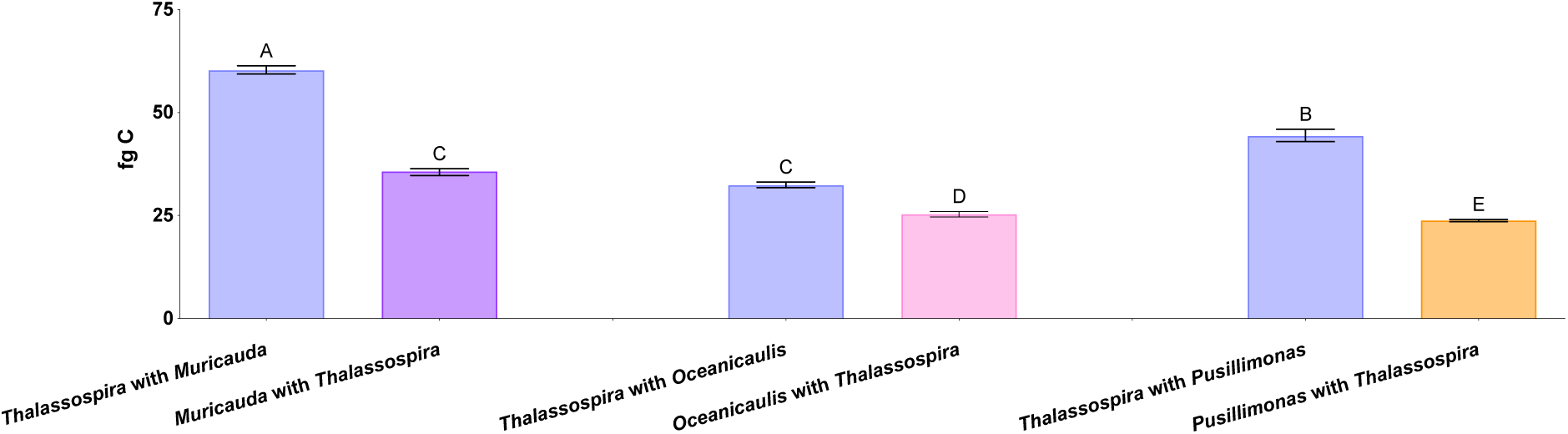
Median single-cell carbon content in the Thalassospira co-cultures.

*Stappia* CUE was reduced with *Muricauda*, similar to *Thalassospira* cultured with *Muricauda*, perhaps also in response to antagonism. Indeed, biomass production rate (Fig. 5B), CO_2_ production rate (Fig. 5B), and median cell C content were all reduced (Fig. S5). These observations were consistent with production of recalcitrant metabolites by *Muricauda* that may be linked to growth reduction in the diatom *Phaeodactylum tricornutum* [68]. Antagonism has been found to decrease CUE in the target organism as C resources are used for defense [69]. At larger scales, these interactions may contribute to the net heterotrophic state of the ocean via faster conversion of DOM to CO_2_ [70, 71].

The similarly reduced CUE for *Stappia* co-cultured with *Pusillimonas* indicated potential antagonism as well, which was notably the opposite response of *Thalassospira*. In addition to reduced biomass and CO_2_ production rates (Fig. 5B), *Stappia* exhibited the lowest median cell carbon content values in this co-culture (due to the smallest cell sizes), alluding to potentially negative shifts from biomass production in the presence of *Pusillimonas*. Certain *Pusillimonas* spp. may exhibit antifungal activity [72, 73], but its antibacterial properties have not been reported in the literature. Notably, as members of the Burkholderiales order, they are related to Burkholderia spp., which release bactericidal compounds effective against a variety of targets [74, 75], including conspecifics [76]. Antagonistic interactions are proposed to occur most intensely in densely populated microenvironments, such as marine organic particles [77], and may shift community composition by affecting sensitive taxa. Our results imply less deleterious relationships may be occurring within free-living bacterial consortia as well, with important consequences for C fate. The deviations from the patterns seen in the *Thalassospira* co-cultures demonstrated how genus-driven interactions altered C metabolism and underscore the complexities of microbial and chemical ecology that underlie biogeochemical cycles.

**Figure S6.**
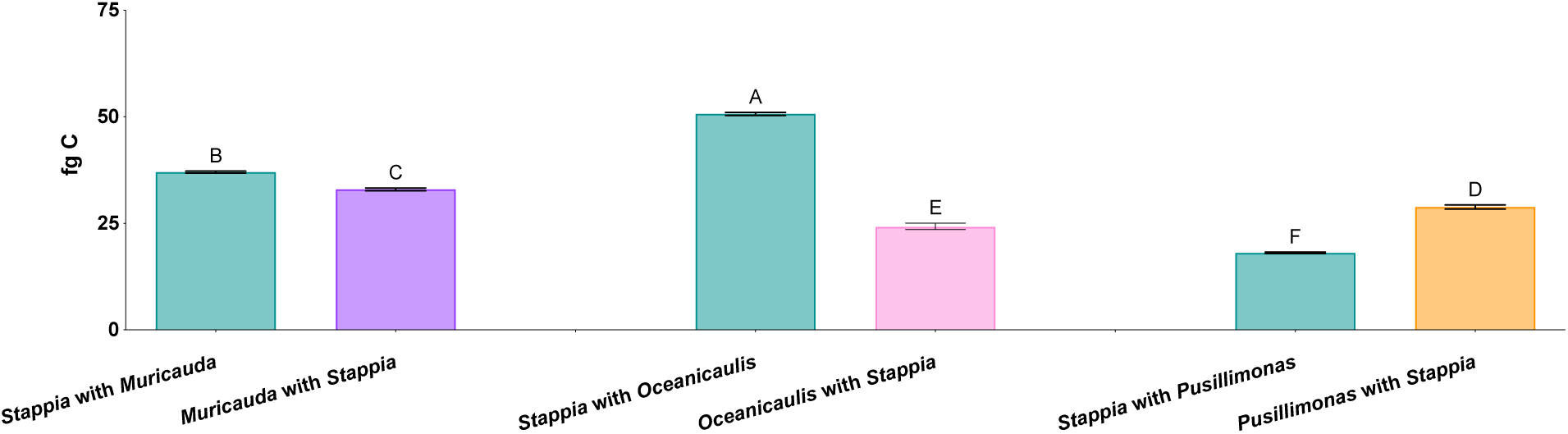
Median single-cell carbon content of Stappia co-cultures

### 3 Non-incorporators exhibited greater CUE of glycolate-derived substrates from *Stappia*, linking ecology to C fate of a major component of marine DOM

In ecological settings, *Muricauda* appears poised to benefit from interactions with *Thalassospira* and *Stappia* as the acquisition of glycolate-derived C would offer a respite from the ‘boom and bust’ dynamics defined by nutrient deficient periods that punctuate algal blooms or high particulate organic matter environments [78, 79]. *Muricauda* populations could be more likely to persist in community partnerships with *Stappia* compared to *Thalassospira*, based on the higher CUE values that were driven by greater anabolism (Fig. S6). These results underscore the influence incorporator and non-incorporator identities have on the fate of glycolate carbon, with higher conversion to biomass and less to CO_2_ in a system dominated by *Stappia* compared to *Thalassospira*.

**Figure S7.**
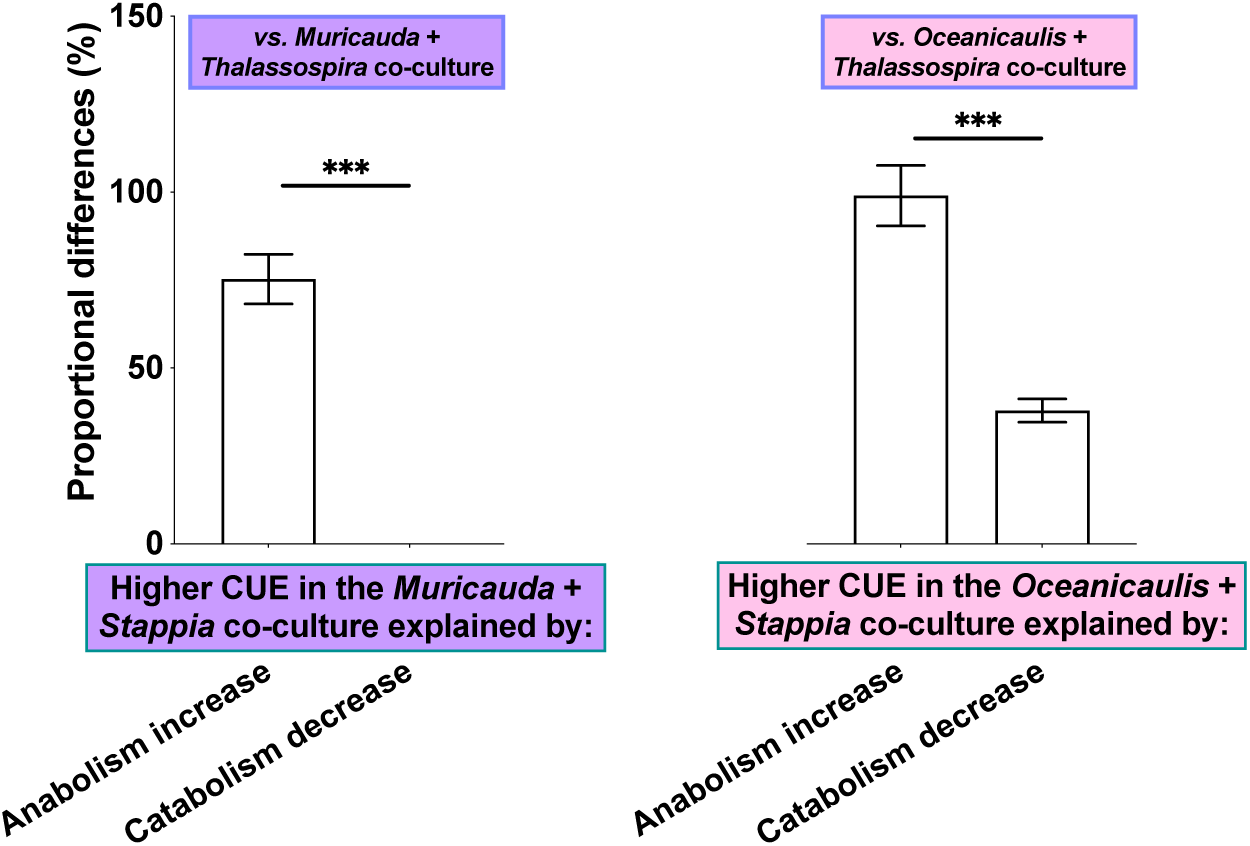
Proportional changes in anabolism (biomass production) versus catabolism (CO_2_ production) driving enhanced CUE in the Muricauda + Stappia (A) and Oceanicaulis + Stappia (B) co-cultures. CUEs of Muricauda and Oceanicaulis are higher when co-cultured with Stappia than Thalassospira due to increased retention of glycolate C into biomass, as opposed to decreased conversion of glycolate into CO_2_.

The influence of incorporator identity on *Oceanicaulis* metabolism of glycolate-derived substrates is likely related to its reliance on compositionally diverse microalgal exudates for growth [80]. Further, the ability of *Oceanicaulis* to metabolize the glycolate-derived compounds may rely on the presence and preferential metabolism of other substrates (i.e., carbon catabolite repression) [81], perhaps specifically requiring dissolved organic nitrogen [40]. Substrates released by *Thalassospira* only partially fulfilled this need for most *Oceanicaulis* cells while substrates from *Stappia* featured characteristics that enabled more uptake in a smaller population of *Oceanicaulis*. These interactions with both *Thalassospira* and *Stappia* mobilized more glycolate-based C to CO_2_ while highlighting how complexities in cross-feeding and other cooperative metabolic interactions affect our mechanistic understanding of substrate fate, even in a simple laboratory system.

*Pusillimonas* exhibited relatively minimal responses to incorporator identity, indicating low suitability of secondary metabolites from both *Thalassospira* and *Stappia* for metabolism. As an efficient incorporator of algal DOM [40], *Pusillimonas* may prefer algal exometabolites to support its anabolic needs [82]. Indeed, algal-bacterium-bacterium relationships influence both metabolism and growth of both interacting bacterial strains [83]. Combining these studies to examine C fate within a tripartite cross-feeding system would offer next level complexity to explore how substrate preferences affect C fate in a more naturalistic system.

## Conclusions and future directions

It is well-established that bacterial metabolism is a major contributor to the net trophic states of ecosystems [71, 84, 85], but how concerted bacterial metabolisms modulate more cryptic and nuanced C transformations is less clear and straightforward to examine. Using glycolate as a sole carbon source in simplified microbial systems, we presented here how the relationship between bacterial taxa and cross-feeding within a consortium exerts control over C fluxes. Our findings indicate that bacterial CUE of glycolate is species-specific and that changes in CUE can be driven by identity of co-occurring taxa. The co-occurring taxa unable to assimilate a specific substrate maintain participation in C cycling of that substrate through metabolism of cross-fed secondary metabolites, which feature secondary CUEs influenced by taxonomy of the primary incorporator. Given a ∼1 Pg C yr⁻¹ glycolate flux globally, the CUE ranges calculated here estimate that cross-feeding would divert, on average, an additional 2.8 Tg C into secondary bacterial production and 81.2 Tg C into secondary CO_2_ production. It is expected that glycolate cycling would occur most intensely in regions of high autotrophic glycolate production and/or high heterotrophic consumption, such as mesotrophic zones [7]. However, the complete drawdown of glycolate by *Thalassospira* and *Stappia* shown here suggests that regions with low glycolate production may reflect rapid consumption. Therefore, developing approaches to quantify glycolate concentrations without the influence of bacterial assimilation will enable a more accurate accounting of glycolate contribution to global CO_2_ production. Melding fieldwork with lab-scale studies will be necessary in upcoming decades as the compounding drivers of high light [86], high temperatures [87], and altered O_2_/CO_2_ ratios [88] may alter algal-bacterial physiology [89] and increase rates of photorespiration and glycolate exudation. These dynamics are also critical to our understanding of the microbial pump carbon (MCP) in the current and future ocean [90]. Cross-feeding networks in microbial communities that shunt more C toward biomass may: 1) increase bacterial stocks that eventually sequester C through production of recalcitrant necromass [91, 92] and/or 2) maintain biomass production that enables continued release of potentially recalcitrant compounds [93, 94]. Importantly, other substrates must be considered as well, not only because they are components of the organic matter pool, but because their compositions impact their eventual fate. The CUE values of glycolate reported in this work agree with the low CUEs of bulk communities reported in other studies, which range 5 to 40% [9, 95], and highlight how the substrate’s reduction potential per unit carbon (ɣ_s_) influences CUE. Glycolate has a ɣ_s_ of 3 while more reduced substrates such as leucine have values of 5 and above. Coupled modeling-experimental works that focus on this interplay between bacterial community composition, substrate reduction potential, cross-feeding, and C fate will be needed to further refine our predictive capabilities of elemental fluxes in aquatic ecosystems.

## Acknowledgments

This work was performed under the auspices of the U.S. Department of Energy by Lawrence Livermore National Laboratory (LLNL) under Contract DE-AC52-07NA27344. Funding was provided by LLNL’s Lab Directed Research and Development (LDRD) Program grant 19-LW-044, release number LLNL-JRNL-2011611. We thank Christine Shulse for constructive comments that improved the manuscript and Christy Ramon for assistance with NanoSIMS sample preparation and logistics.

## Data Availability

Source data file is available in the supplementary information. Code for bioinformatics analysis is available on github (https://github.com/jeffkimbrel). Bacterial isolates available upon email request to.

## Materials & Methods

### Comparative genomics of glycolate metabolism and global distribution of isolates

All isolates in this work were obtained from initially complex consortia associated with the diatom *Phaeodactylum tricornutum* [21]. KEGG orthologs (KOs) were identified by scanning the isolate genomes with KofamScan v1.3.0 [97] with hidden markov models of the KEGG 11336689 release [98] using the model-defined trusted cut-offs. Glycolate metabolism and related KOs were identified as follows: phosphoglycolate phosphatase, K01091; glcA, K02550; glcB, K01638; glcC, K11474; glcD, K00104; glcE, K11472; glcF, K11473; isocitrate lyase, K01637; lactate permease, K03303/K00427; BHAC, K18425. Global abundances of taxa closely-related to the isolates was determined by comparing the 16S rRNA gene sequences from the isolate genomes against the Ocean Gene Atlas (OGA)[34, 35]. The OGA has prokaryotic sequences from both the Tara Oceans [99]and Global Ocean Sampling [100] datasets.

### Experimental cultivation conditions to examine exchange of glycolate-derived compounds between incorporators and non-incorporators

We used a co-culturing platform (Fig. S1) that enables side-by-side culturing of microbes separated by a 0.1 µm pore size membrane. The goal for this work was to examine the influence of microbial identity on carbon fate by comparing ^13^C-labeling of single cells and bulk glycolate-derived CO_2_ between incorporators alongside non-incorporators. The primary mechanism hypothesized to be in action was the release of glycolate-derived extracellular metabolites from the incorporator (i.e., ^13^C-labeled secondary substrates) that were subsequently assimilated by the non-incorporator.

Isolates were cultivated in ZoBell media (5 g peptone + 0.5 g yeast extract diluted and autoclaved in artificial seawater) for six days at room temperature RT. Turbid cultures in late stationary phase were generated during this period, as noted by daily measurements of OD_600_. The cultures were then diluted 100-fold into fresh, 100-fold diluted Zobell media (Z/100) for a 24 h incubation. The cultures were then washed 3x with artificial seawater media via centrifugation at 16,000 × g and resuspended in the same volume of artificial seawater containing 50 µM ^13^C-glycolate (99 atom% ^13^C; Cambridge Isotopes Laboratories Inc., Cambridge, MA, USA), 13 µM ammonium chloride (NH_4_Cl), and 1.4 µM monosodium phosphate (NaH_2_PO_4_). Assuming starting concentrations of 10^8^ – 10^9^ cells mL^-1^ in ZoBell media containing 367.5 mM C (based on estimates of C content in peptone and yeast extract from manufacturer specifications), this stepdown of cell and nutrient concentrations established 10^6^ – 10^7^ cells mL^-1^ in liquid containing approximately 3.7 mM C in Z/100 followed by 100 µM C in the glycolate media.

The first experiment consisted of incubating the above-described monocultures for 42 h in autoclaved sealed serum vials and then proceeding with the analyses described below. The second set of experiments examined metabolism and cross-feeding of glycolate in co-cultures consisting of a glycolate incorporator and a non-incorporator. Samples were prepared Identically as the first monoculture experiment and then cultivated in duplicate gas-tight autoclaved co-culture vessels (Fig. S1; custom version of MFC500.40.2, Adams-Chittendens Scientific, Berkeley, CA) with the glycolate incorporator on one side and the non-incorporator on the other separated by a 47 mm diameter 0.1 µm polycarbonate filter (VCTP04700, Millipore Isopore). Co-cultures were incubated for one week and then sampled for analyses described below.

### Microscopy and image analyses to calculate cell concentrations and biomass carbon

Sterile needle syringes were used to pierce septa and acquire subsamples (1 – 10 mL) that were then fixed with 37% formaldehyde (3.7% final concentration, 0.2 µm filtered) for 1 hour and filtered onto 0.2 μm pore size white polycarbonate membrane filters (Whatman Nuclepore, GE Healthcare Life Sciences, Pittsburgh, PA). After washing and drying, filters were stored at -20°C until analysis. Filters were cut with sterile scissors into 1/8 slices, and three to four pieces were placed onto ethanol-cleansed microscope slides. Mounting and staining were completed in a single step using an antifade + staining mounting medium prepared in 0.2 µm filtered 1:1 PBS:glycerol containing 0.1% *p*-phenylenediamine and 1 µg mL^-1^ 4ʹ,6-diamidino-2-phenylindole (DAPI). Thirteen microliter droplets were pipetted onto 24x50mm #1.5 coverslips with locations that corresponded to those of the filter pieces. The coverslips were then inverted on top of the slides to sandwich the filter pieces for 10 – 15 minutes of stain time, after which bubbles and excess mounting medium were gently pressed out. A minimum of ten images were acquired on a Leica DMI6000b widefield fluorescence microscope and exported as tagged image file formats (tiffs). A custom sci-kit image [101] v0.17.2 Python script was applied to obtain ROI dimensions using watershed segmentation on a local threshold mask (https://github.com/jeffkimbrel/jak-bio). Length (l) and width (w) dimensions were converted to biovolumes using the equation:

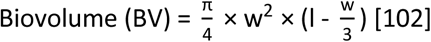

Biovolumes were then converted to carbon content with the equation:

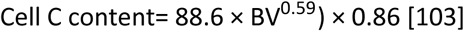

**Figure S46.**
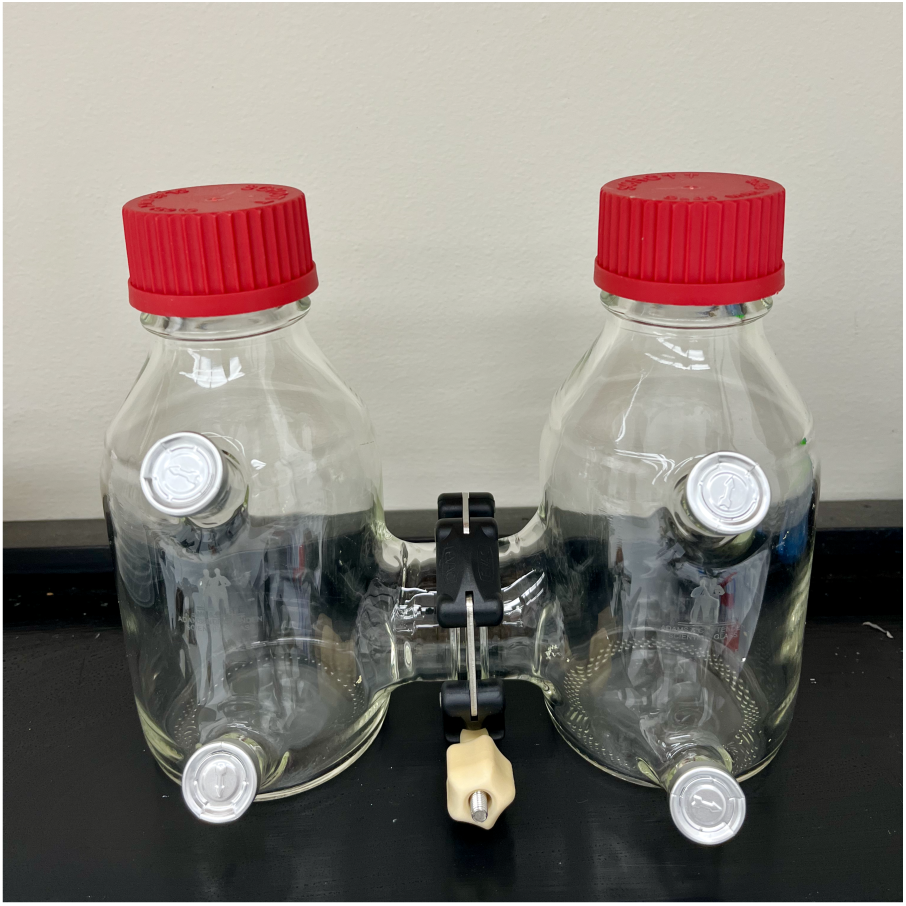
Co-culture vessel used in this study featuring ports for liquid and headspace sampling.

### Nanoscale stable isotope probing to measure ^13^C-glycolate incorporation

NanoSIP analyses [104] were performed on single cultures in the presence of labeled glycolate to quantify glycolate-specific C incorporation and glycolate-derived cross-feeding of C compounds from glycolate incorporators to non-incorporators – all at the single cell level. Wedges of 1/8 size were cut out of the same polycarbonate filters used for microscopy with sterile scissors and adhered to an aluminum electron microscopy stub using conductive tabs (#16084-6, Ted Pella, Redding, CA) and sputter coated with ∼5 nm of gold. Isotope imaging was performed with a CAMECA NanoSIMS 50 at Lawrence Livermore National Laboratory. The 16 keV primary ^133^Cs+ ion beam was set to 2 pA, producing an approximately 150 nm diameter beam. Rastering was performed over 20 x 20 μm analysis areas with a dwell time of 1 ms pixel^-1^ for 19-30 scans (cycles) and generated images containing 256 x 256 pixels.

Sputtering equilibrium at each analyses area was achieved with an initial beam current of 90 pA to a depth of ∼60 nm, thus ensuring analysis of intracellular isotopic material. After tuning the secondary ion mass spectrometer for mass resolving power of ∼7000 (after 1.5x correction), secondary electron images and quantitative secondary ion images were simultaneously collected for ^12^C_2-_, ^13^C^12^C^-^, and ^12^C^14^N^-^ on individual electron multipliers in pulse counting mode, as outlined in [104]. All NanoSIMS datasets were initially processed using L’Image (http://limagesoftware.net) to perform deadtime and image shift corrections of ion image data before extracting ^13^C^12^C^-^/^12^C_2-_ ratio data, which reflected the level of ^13^C incorporation into biomass from ^13^C-glycolate (^13^C/^12^C = 0.5 × ^13^C^12^C^-^/^12^C_2-_). Regions of interest (ROIs) for isotopic ratio quantification were drawn individually around each cell using the L’image particle analysis function using the ^12^C^14^N^-^ images and exported for statistical analyses in GraphPad Prism. Isotope incorporation data for cells were normalized to the ^13^C enrichment detected on the polycarbonate filter for each sample, to enable cross sample comparisons. Data are reported as a Cnet rate, which is the percent of biomass produced from 13C-glycolate per day as calculated in [41].

### Cavity ring down spectroscopy and gas chromatograph to measure ^13^C-glycolate respiration to ^13^CO2

After sample collection for NanoSIMS and microscopy, the remaining media was acidified with 4.5 N sulfuric acid (H_2_SO_4_; 281.25 nM final concentration per Casey et al. 2017 [13] to convert dissolved inorganic carbon to CO_2_ in the headspace over the span of two days. Ten milliliters of headspace volume was sampled and transferred to sterile 20 mL evacuated serum vials. Using a Picarro G2201-I cavity ring down spectrometer (CRDS), 5 mL of sample was removed using a gas-tight syringe with stopcock and injected into the small-sample isotope module (SSIM) port. The factory-defined run routine was started, which calibrates to an internal standard prior to sample analysis. The ^13^CO_2_ measurements (in units of parts per million; ppm) were dilution-corrected using an equation from the manufacturer. It multiplies the ^13^CO_2_ concentration by the ratio of the zero-air pressure (O_2_ + N_2_, no CO_2_; user selected 5 - 10 psi) that fills and maintains the required 20 mL of gas in the SSIM chamber to the pressure of the sample within the chamber. Important details include: 1) multiplying psi by 51.7 to convert to Torr and 2) correctly calculating total zero air pressure by adding atmospheric pressure (psia; 14.7) to the zero-air pressure coming from the tank (set to 7.5 psi for this work). We include these calculations below for clarity:

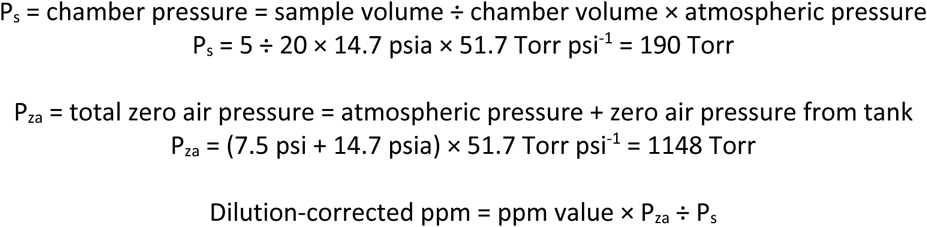

### Calculation of glycolate ^13^C-carbon comprising microbial biomass and CO2 (Cnet)

^13^C-enrichment data (^13^C/^12^C ratios) for cells and headspace generated form the nanoSIMS and CRDS analyses, respectively, were used to calculate the percent of isotope in cells or headspace along with background and substrate (glycolate in this case) ratios normalized to incubation time to generate a rate. As described previously [41, 105], this is represented by:

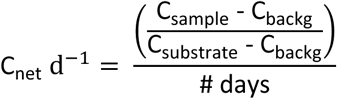

### Derivatization and GC-MS analysis to detect remaining glycolate in the media

We modified an existing method to derivatize and detect glycolate using GC-MS [106]. Three milliliters of 0.2 µm filtered culture extracts were placed in glass vials and dried under a stream of nitrogen gas at 50°C for 15 minutes. Once the samples were dry, 50 µL of pyridine followed by 50 µL of BSTFA (*N*,*O*-Bis(trimethylsilyl)trifluoroacetamide) + 1% TMCS (trimethylsilyl chloride) reagent were added to the glass vial and sealed with a Teflon lined septum. The samples were then heated at 60°C for 30 minutes and allowed to cool. An additional 100 µL of dichloromethane (DCM) was added to the sample for extraction, and 100 µL was transferred into a GC-MS vial. The derivatized samples were then analyzed using an Agilent 8890 GC coupled to a 7000D MS/MS mass spectrometer with a PAL-3 autosampler. After multiple syringe washes with DCM, 1 µL of sample was injected into a split/splitless liner with a 10:1 (He:Sample) split flow at 250°C. An Agilent HP5MS-UI column was used for analysis and held at 60°C for 1 minute, then ramped at 10°C/min and held at 315°C for 10 minutes. Glycolate standards were analyzed in scan mode using MS1 to determine the correct m/z for selective ion monitoring (SIM) to improve detection limits. The SIM method used MS1 to select for m/z 205, 147 and 73 ions and used a 30 ms dwell time.

